# Can social media serve as a potential citizen science source for bird-window collision (BWC) data? A study using a decadal data set in Taiwan

**DOI:** 10.1101/2024.03.29.587372

**Authors:** Chi-Heng Hsieh, Gen-Chang Hsu, Ling-Min Wang

## Abstract

Citizen science is increasingly used in bird-window collision (BWC) research to collect data. However, few studies have collected BWC data from social media, and it remains unknown whether the data quality is comparable to that of reported on dedicated platforms. To evaluate the potential of social media as a citizen science data source for BWC, we collected BWC data on social media Facebook as well as Taiwan Roadkill Observation Network (TaiRON), the main dedicated citizen science platform for reporting wildlife mortalities in Taiwan. We compared a decade of BWC data (2012–2022) from the two platforms by examining the nationwide geographical coverage and the species compositions of the BWC observations. Overall, we recorded 2,583 BWC cases involving 153 BWC species from Facebook, and 1,000 BWC cases involving 104 BWC species from TaiRON. More than half of the BWC individuals from Facebook were not found dead when observed, whereas all records on TaiRON were dead individuals. The nationwide geographical coverage and the species compositions of the top 80% cumulative BWC individuals were generally similar between the two platforms. Moreover, the sampling completeness of the two platforms both exceeded 95% (Facebook: 98.0%; TaiRON: 96.0%). To our knowledge, this study is among the first to collect BWC data through social media posts, and our results show that the quantity and quality of Facebook data can be comparable to that of the well-developed citizen science platform TaiRON. Taken together, social media Facebook may not only serve as a promising tool for collecting BWC data, but also provide a platform for public education, which can benefit bird conservation. Finally, integrating data from different citizen science sources helps paint a more complete picture of BWC patterns, especially in understudied areas such as Asia.

## 1. Introduction

Bird-window collision (BWC) has been an increasingly important conservation issue worldwide. Past BWC studies have mostly been conducted in the temperate regions, especially in North America (Loss et al. 2014, Cusa et al. 2015, Hager et al. 2017). However, tropical and subtropical regions may exhibit BWC patterns that are different from those in the temperate regions (Gómez-Martínez et al. 2019, Menacho-Odio et al. 2019, Tan et al. 2023), and there have been calls for more tropical and subtropical studies to address the geographical imbalance in BWC research (Basilio et al. 2020).

Surveying carcasses around human structures is the most common approach to studying BWC. This method provides useful information about the BWC patterns and allows for the estimation of BWC rates and annual cases (Hager et al. 2013, Loss et al. 2014, Hager et al. 2017). However, because of the environmental (warm and humid weather), ecological (abundant scavengers) and cultural (aversion to handling carcasses and frequent cleanups by janitors) differences, the conventional protocol for surveying collided birds (Hager and Cosentino 2014) may not work well in tropical and subtropical countries, which have high carcass removal rates. Besides these limitations, carcass surveying may not be able to accumulate sufficient data to address the emerging BWC issue at a national scale as the approach is both time- and labor-consuming. This is especially true in Taiwan, a subtropical Asian country where BWC research has recently started, and abundant data are needed to understand the BWC patterns.

To overcome these challenges, citizen-science-based methods have been suggested as a powerful tool for engaging the general public in recording large numbers of observations in a relatively short period of time (Cohn 2008, Silvertown 2009, Bonney et al. 2014). In recent years, several citizen science projects on BWC research have been launched to collect abundant BWC data (Bayne et al. 2012, Kummer and Bayne 2015, Loss et al. 2015, Loss et al. 2023). These projects are mainly run on dedicated citizen science platforms, for instance, the Global Bird Collision Mapper (GBCM) (https://www.birdmapper.org/) for Fatal Light Awareness Program (FLAP) Canada and iNaturalist (https://www.inaturalist.org/) for several studies in the United States (Winton et al. 2018, Brown et al. 2019). In Taiwan, the Taiwan Roadkill Observation Network (TaiRON) is the main dedicated collaborative citizen science platform for documenting wildlife mortalities. Launched in 2011, this platform provides a standard guideline for observers to report various types of wildlife death incidents in Taiwan (e.g., roadkill, bird-window collisions, intoxication, and animal-caused mortalities) (Chuang et al. 2016).

In addition to the dedicated platform such as TaiRON, social media can be a potential source for gathering citizen science data (Newman et al. 2012, Oliveira et al. 2021, O’Neill et al. 2023). Facebook (Facebook), one of the most widely used social media in Taiwan (around 19 million active accounts out of a population of 23 million people as of 2023), may act as a good pair of “eyes” in search for abundant BWC data. The flexibility in the formats and contents of Facebook posts enable users to share more information about the BWC observations (e.g., how they dealt with the collided birds, what the states of the collided birds were, and what they thought about BWCs etc.). Moreover, similar to recall surveys, which are increasingly common in BWC research (Bayne et al. 2012, Kummer et al. 2016, Rebolo-Ifrán et al. 2019, Żmihorski et al. 2022), Facebook provides access to relevant public posts through keyword search, allowing researchers to retrieve BWC observations posted in the past years.

Despite the aforementioned advantages, social media have rarely been used in past BWC studies as a data source. It remains unknown whether social media can be used to collect BWC data, how much time and effort is required, and whether the data quality is comparable to that of dedicated platforms. To fill these gaps, we first collected BWC data on both Facebook and TaiRON. We then compared the similarity and difference between the two platforms by examining the nationwide geographical coverage and the species compositions of the BWC observations from the two data sources. Additionally, we quantified the sampling completeness of the two data sources to assess the adequacy of data collection efforts. Our aims are to evaluate the potential of social media (Facebook) as a citizen science data source for BWC research, and to better understand BWC patterns in less-studied regions (Taiwan).

## 2. Materials and Methods

### 2.1 Collection of BWC data from Facebook and TaiRON

In March 2019, Raptor Research Group of Taiwan initiated a public Facebook group “Reports on Bird-Glass Collisions” (https://www.facebook.com/groups/birdwindowcollision), which encourages members to post their own observations or share others’ observations, distribute relevant information, and discuss topics on BWCs. The posts in the group were checked regularly, and if a contributor posted or shared a BWC observation, the person was contacted for more information on the date of observation, location (GPS coordinates if possible), species, number of individuals, simple descriptions of the observation, and photos if taken. We documented these information as the first part of our Facebook BWC data from March 2019 to December 2022.

Considering there might be BWC observations posted on Facebook before the group “Reports on Bird-Glass Collisions” was initiated or by users who were not in the group, between February 2020 and December 2022, we carried out systematic searches through public Facebook posts on a weekly basis using a series of search terms “bird AND collide” and “bird AND collide AND (glass OR window OR door OR wall)”. The keyword “bird” was replaced by the common names for certain families of birds (Accipitriformes, Anseriformes, Galliformes, Passeriformes, and Strigiformes) once a month in case the contributors used the common names instead of “bird” *per se* in their posts. Additionally, in March 2022 and December 2022, systematic censuses were conducted with the keyword “bird” replaced with the common names of all birds on the 2020 Checklist of the Birds of Taiwan (Ding et al. 2020). All systematic searches and censuses were carried out in Taiwanese Mandarin (see Appendix Methods for more details on the search queries). The BWC observations collected via these backtracking methods comprised the second part of our Facebook BWC data. (These BWC posts were also shared to the “Reports on Bird-Glass Collisions” Facebook group.)

To collect BWC observations from TaiRON, we requested data on avian records of which the cause of death was reported as “window collisions” from the platform manager in December, 2022. Only complete reports (i.e., containing information on the date of observation, location [GPS coordinates], species name, and photos of the bird) were retained as the final TaiRON BWC data. There were about 100 duplicate observations reported on both Facebook and TaiRON. For these cases, the observations on TaiRON were retained since TaiRON is a well-developed citizen science platform with standardized entry formats. Moreover, some observations contained multiple collided birds. Hereafter, we referred to “BWC case” as a single observation (a Facebook post or a TaiRON record) and “BWC individual” as a single collided bird. Escaped pet species without a wild population in Taiwan (e.g., parrots) were excluded from the BWC data.

To ensure that the species information from the contributors on Facebook and TaiRON was correct, we further identified the BWC individuals (to species level if possible) using the provided photos. While all cases reported on TaiRON were found dead, cases reported on Facebook were not limited to dead or live individuals. Based on the photos and the descriptions by the contributors, the non-dead BWC individuals on Facebook posts were further classified into four different states (Fig. 1): “Flew away”, “Sent to rescue center”, “Captured”, and “Uncertain”. “Flew away” refers to individuals that left the spot by themselves after the BWC; “Sent to rescue center” refers to individuals that were sent to rescue centers by the observers; “Captured” refers to individuals (often stunned or severely injured) that were restrained by human; “Uncertain” refers to BWC individuals whose states couldn’t be clearly defined from the photos or the descriptions.

**Figure 1.**
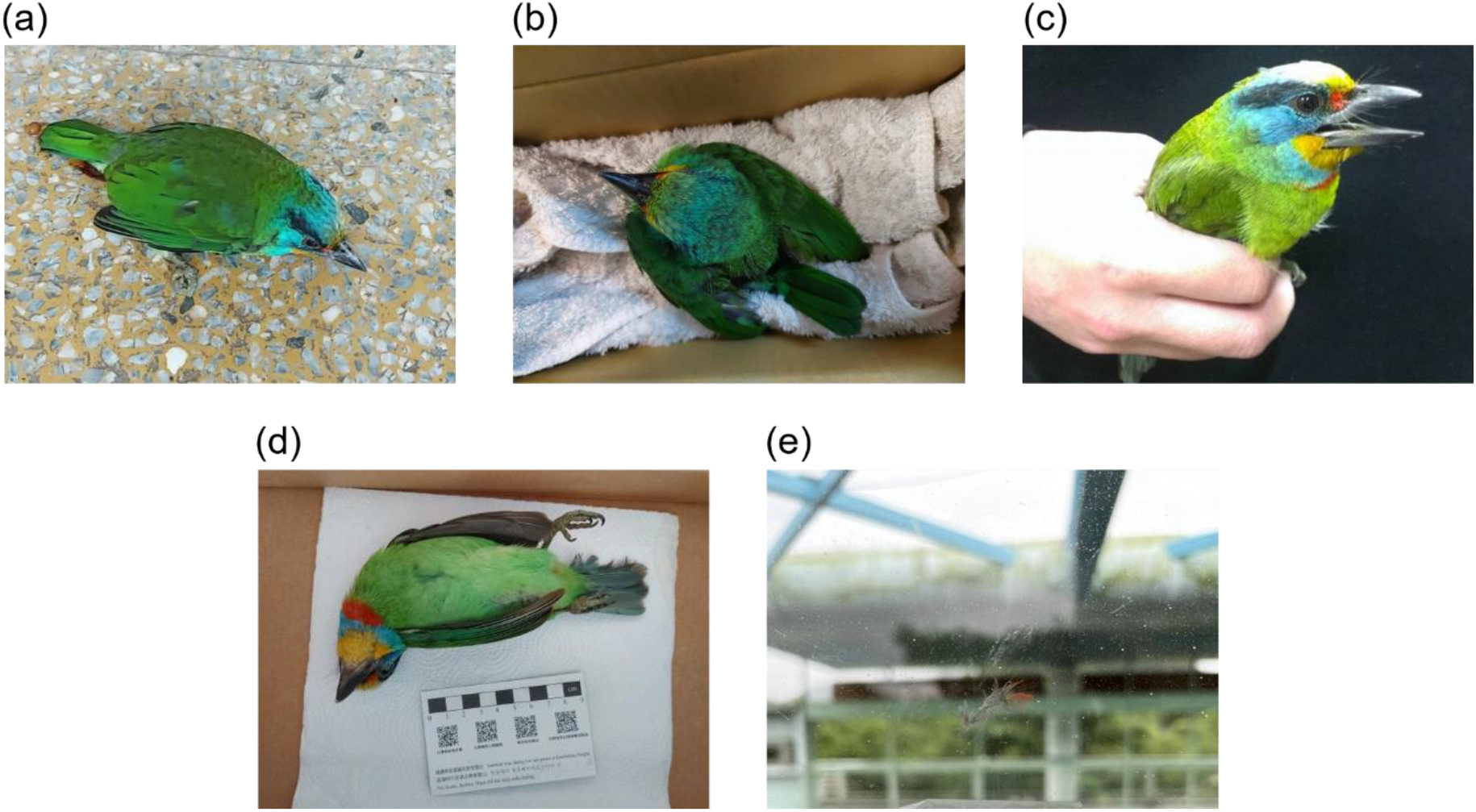
Example photos of different states of collision individuals (Taiwan Barbet *Psilopogon nuchalis*) on Facebook posts: (a) Flew away (the individual was found stunned and flew away by itself later); (b) Sent to rescue center; (c) Captured; (d) Found dead; (e) Uncertain (only feathers left on the glass). See *Materials and Methods* for more details on the state classification. Photo credits: Si-Min Lin (c); Chia-Yun Kan (e).

### 2.2 Data analysis

The BWC records dated back to 1992 for Facebook and 2005 for TaiRON (Fig. 2). To ensure that the data periods were comparable between the two platforms and considering that there were only six BWC records from TaiRON before 2012, we therefore restricted our comparisons to data between 2012 and 2022. Individuals not identified to species level were excluded, resulting in a total of 3,550 BWC individuals (94.5% of the 3,754 BWC individuals in the original full data set) in the final data were analyzed.

**Figure 2.**
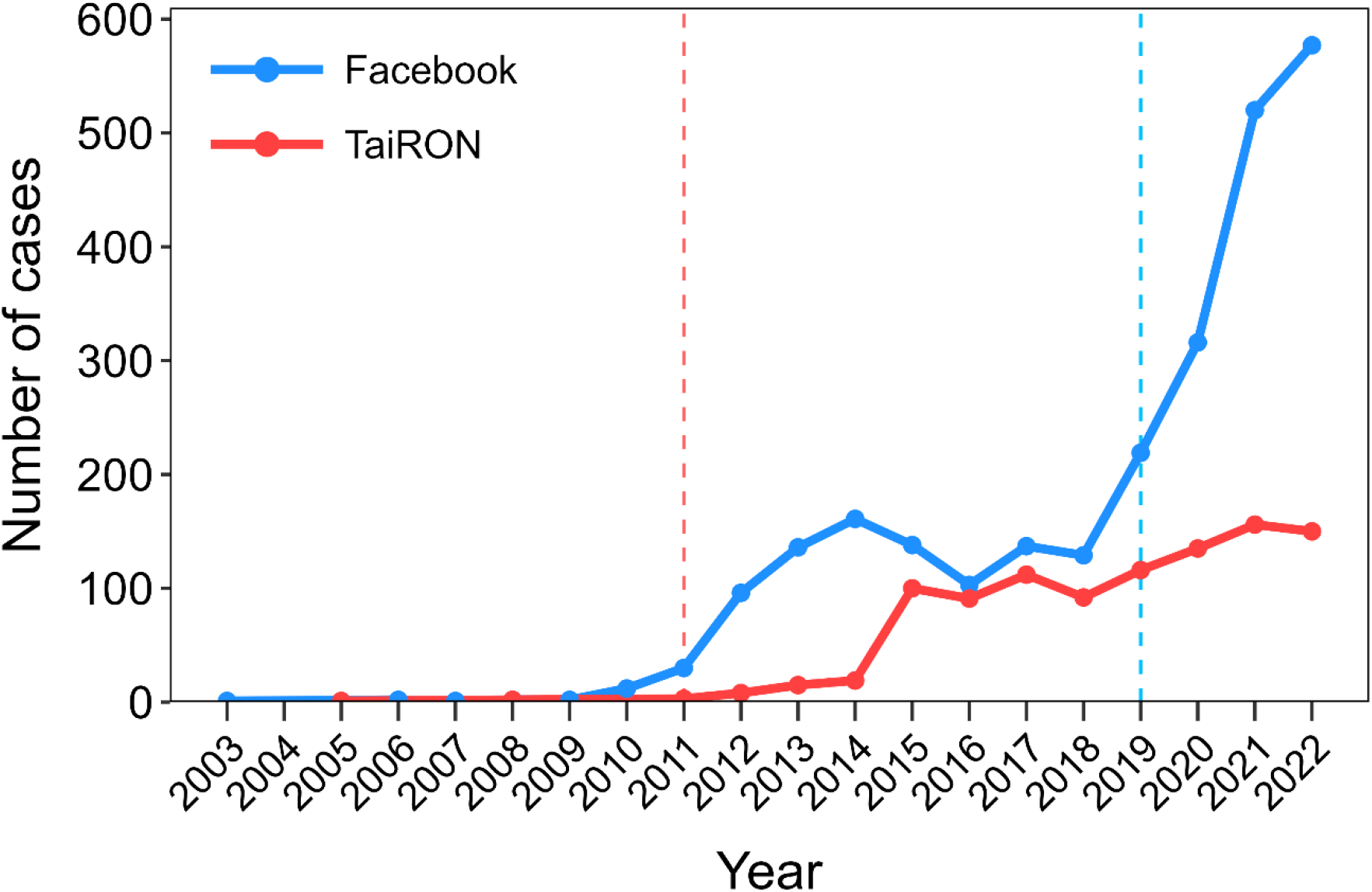
The number of collision cases collected from Facebook and TaiRON over years. Dashed lines denote the start of the collision data collection on the two platforms. Note that one collision case in 1992 collected from Facebook was excluded from the figure to improve visualization.

To compare the rank-abundance distributions and species patterns of the BWC data collected from Facebook and TaiRON, we constructed the rank-abundance curves based on the proportion of each BWC species and examined the species compositions of the 80% cumulative BWC individuals from the two platforms. To determine the number of BWC species and sampling completeness of the BWC data from Facebook and TaiRON, we constructed sample-size-based interpolation and extrapolation curves using the R “iNEXT” package (Hsieh et al. 2016). The rarefaction estimates and the associated 95% confidence intervals were computed based on 1000 bootstrap samples. The number of BWC species and sampling completeness of the two platforms were compared at the reference point rarefied to the same number of BWC individuals from TaiRON (which had the lower number of BWC individuals of the two platforms). All analyses were performed in R version 4.2.1 (R Core Team 2022).

## 3. Results

### 3.1 Data overview

In total, there were 3,583 BWC cases (Facebook: 2,583; TaiRON: 1,000) with 3,754 BWC individuals (Facebook: 2737; TaiRON: 1017) collected from the two platforms. The number of BWC cases increased substantially in recent years (Fig. 2), with the earliest one dating back to 1992 on Facebook.

The BWC cases between 2012 and 2022 from Facebook and TaiRON were mainly located in lowland areas in Taiwan Main Island, while some cases were recorded in the Central Mountain Range and in all outlying islands (Fig. 3a–c). The reported BWC hotspots were concentrated in major cities (e.g., Taipei City, Taichung City, and Kaohsiung City) (Fig. 3d–f).

**Figure 3.**
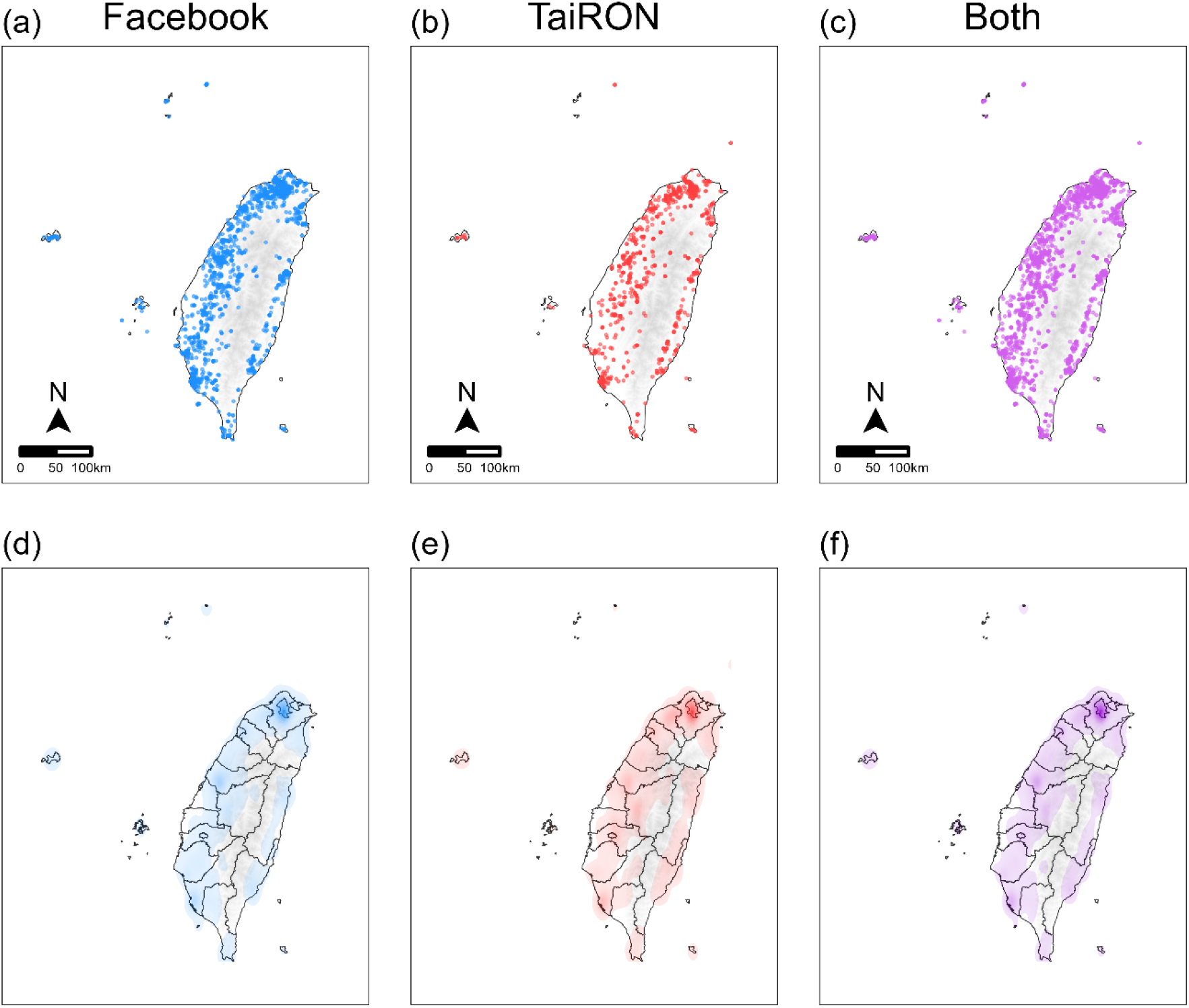
Maps of collision cases between 2012 and 2022 collected from Facebook, TaiRON, and both platforms combined. Panel (a) to (c) show the locations of bird-window collisions with each dot representing a collision case; panel (d) to (f) show the kernel density estimates of collision cases. Note that collision cases without exact GPS coordinates were excluded from the maps.

Of the BWC individuals between 2012 and 2022 collected from Facebook, 41.5% were found dead, 26.8% were captured in the hands of the observers (indicating that the birds were stunned or not able to flee at the moment), 11.6% were sent to the rescue center, 10.3% flew away themselves, and 9.8% could not have their states determined (Fig. 4). All BWC individuals from TaiRON were found dead (Fig. 4).

**Figure 4.**
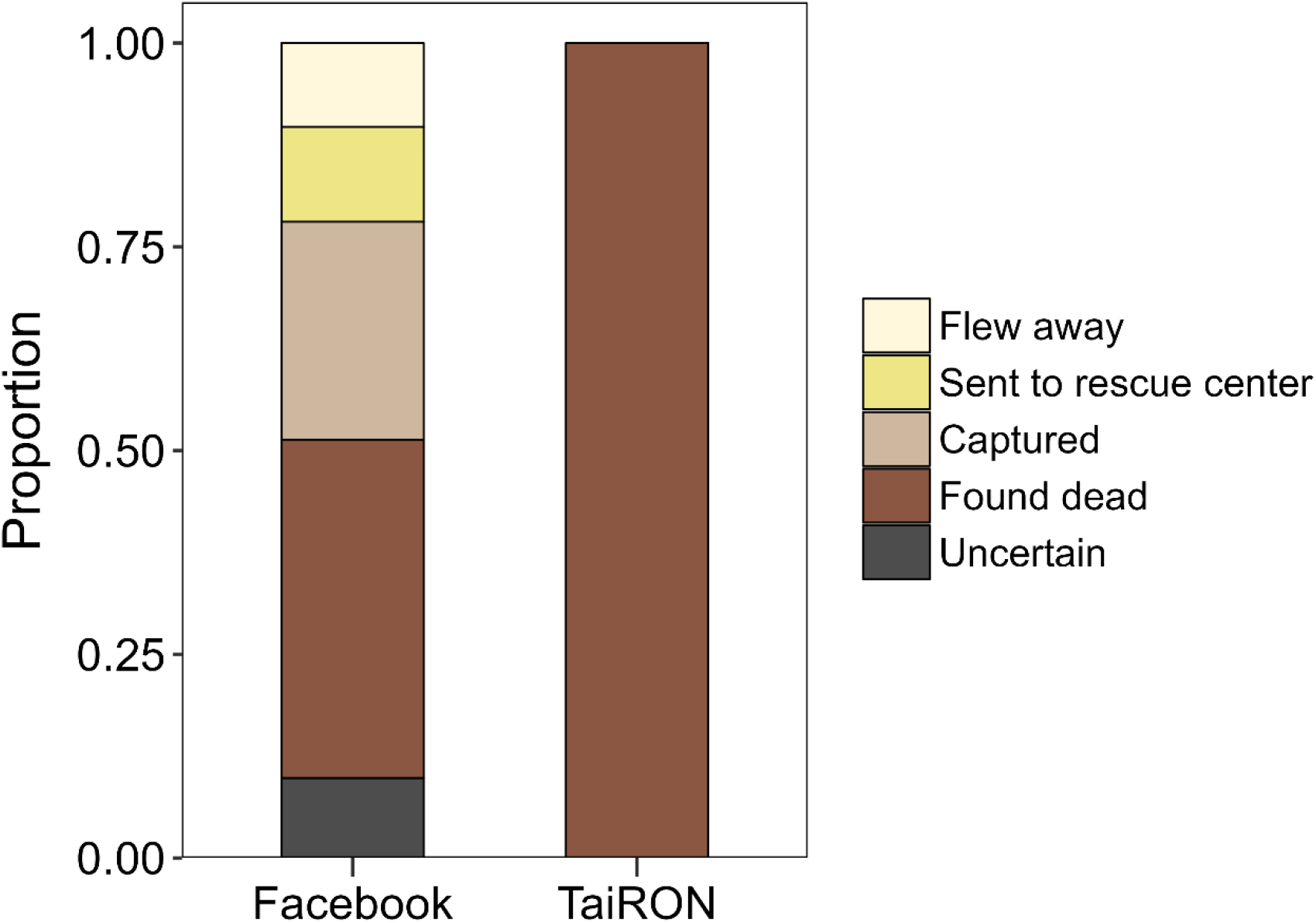
Composition of the states of the collision individuals between 2012 and 2022 collected from Facebook and TaiRON. Note that collision individuals not identified to the species level were excluded from the calculation.

### 3.2 Species patterns of the two platforms

The BWC data from Facebook and TaiRON consisted of 153 and 104 BWC species, respectively (172 species in total when both platforms were combined). There were 85 species present on both platforms; 68 species were present only on Facebook; 19 species were present only on TaiRON (Fig. 5). The rank-abundance distributions of the BWC species were similar between the two platforms, as indicated by the slopes of the curves. Both platforms had a few species with relatively high proportions of BWC individuals (Fig. 6).

**Figure 5.**
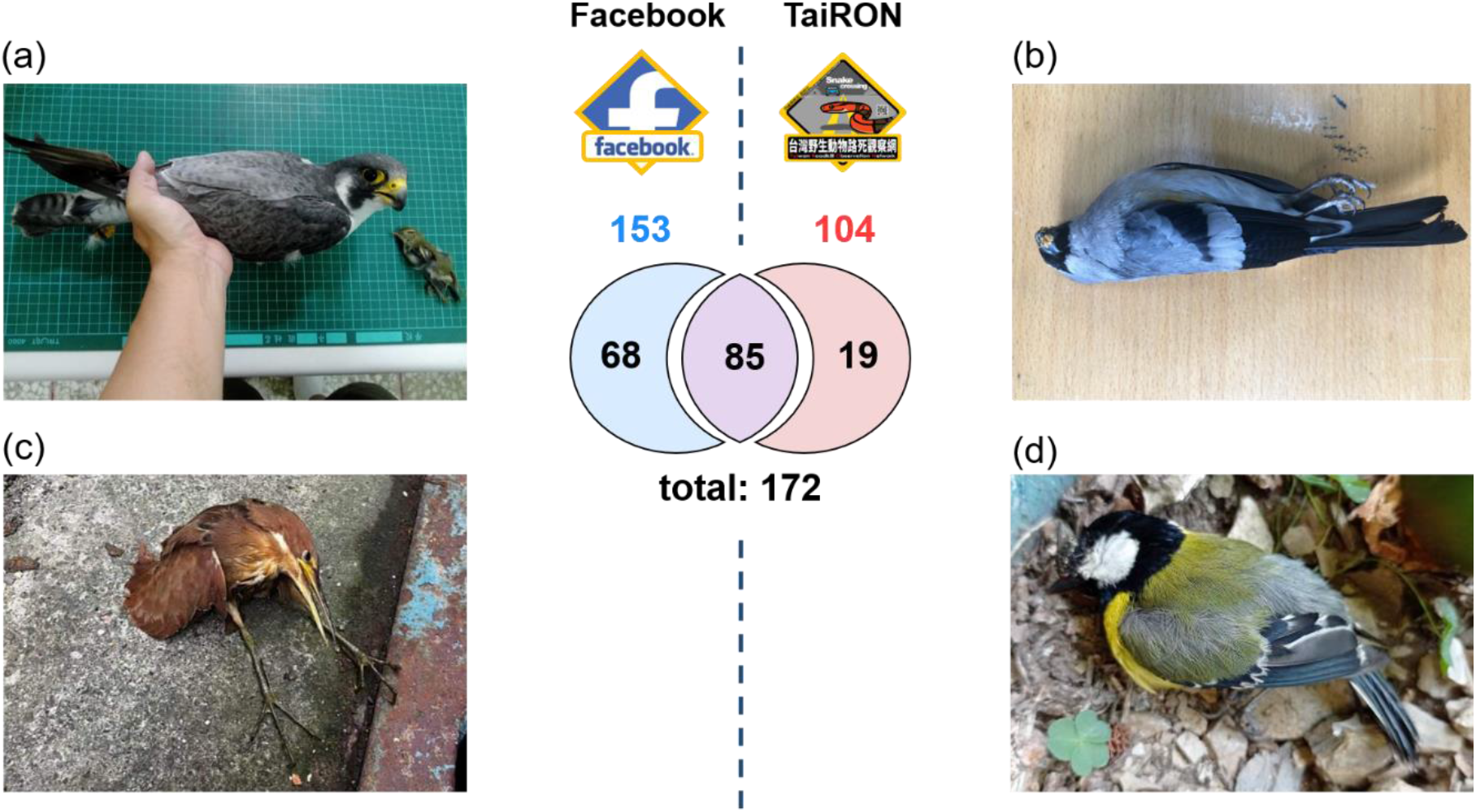
Venn diagram showing the number of collision species recorded on Facebook, TaiRON, and both platforms along with example photos of the species recorded only on Facebook (a), (c) and on TaiRON (b), (d), respectively. (a) Peregrine Falcon (*Falco peregrinus*) (the falcon chased after an Arctic Warbler (*Phylloscopus borealis*) and both collided); (b) Gray-headed Bullfinch (*Pyrrhula erythaca*); (c) Cinnamon Bittern (*Ixobrychus cinnamomeus*); (d) Green-backed Tit (*Parus monticolus*). Photo credits: Taichung Wildlife Conservation Group (a); Yi-Fan Ku (b); Chih-Ping Chang (c); Kuo-Chang Lin (d).

**Figure 6.**
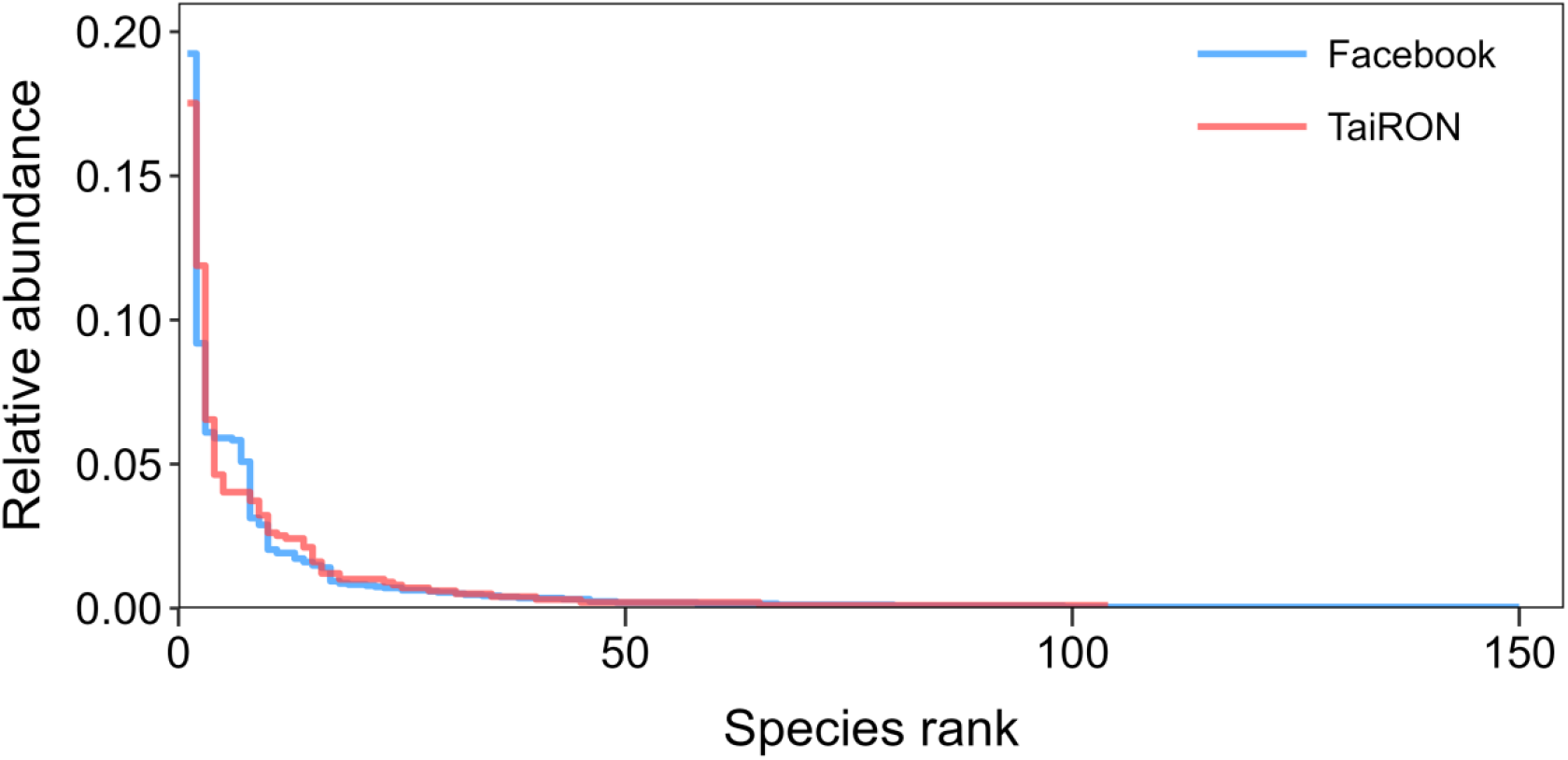
Rank-abundance distributions of collision species in the data between 2012 and 2022 collected from Facebook and TaiRON. The relative abundance of species is computed as the number of collision individuals divided by the total number of collision individuals on each platform. Note that collision individuals not identified to the species level were excluded from the calculation.

The 80% cumulative BWC individuals in Facebook data consisted of 22 species belonging to 13 families: five in Columbidae, three in Turdidae, two each in Accipitridae, Alcedinidae, and Pycnonotidae, and one each in Cuculidae, Estrildidae, Laniidae, Megalaimidae, Passeridae, Strigidae, Sturnidae, and Zosteropidae (Table 1). The 80% cumulative BWC individuals in TaiRON data also consisted of 22 species belonging to 13 families but had a different composition than Facebook data: six in Columbidae, three in Turdidae, two each in Alcedinidae and Pycnonotidae, and one each in Accipitridae, Corvidae, Estrildidae, Hirundinidae, Megalaimidae), Muscicapidae, Passeridae, Sturnidae, and Zosteropidae (Table 1). The majority of species of the 80% cumulative BWC individuals on Facebook and TaiRON were shared; there were four species present only on Facebook (Besra *Accipiter virgatus*, Brown Shrike *Lanius cristatus*, Northern Boobook *Ninox japonica*, and Oriental Cuckoo *Cuculus optatus*) and four present only on TaiRON (White-tailed Robin *Myiomela leucura*, Gray Treepie *Dendrocitta formosae,* Barn Swallow *Hirundo rustica*, and White-bellied Green-pigeon *Treron sieboldii*) (Table 1).

**Table 1.**
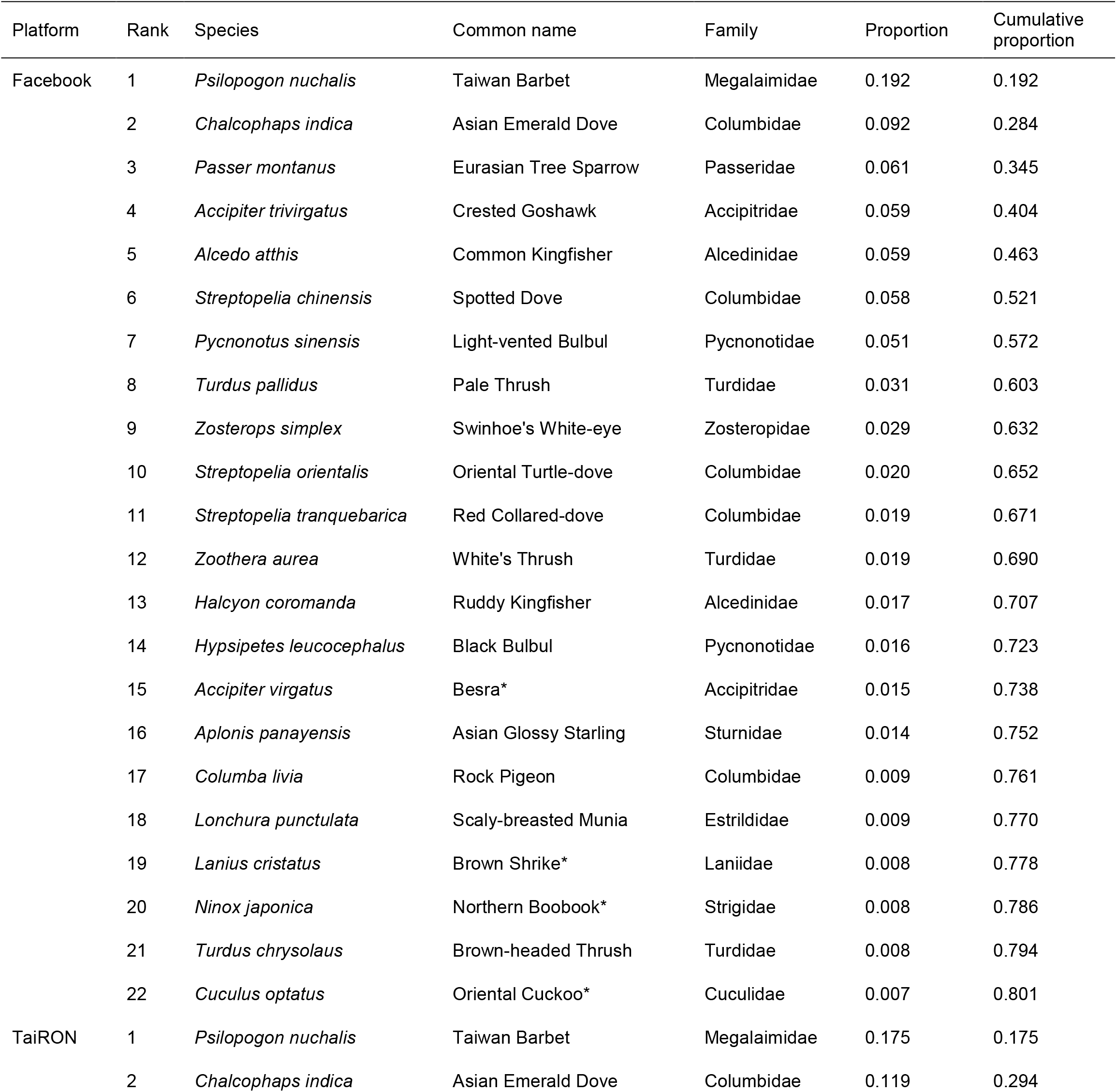
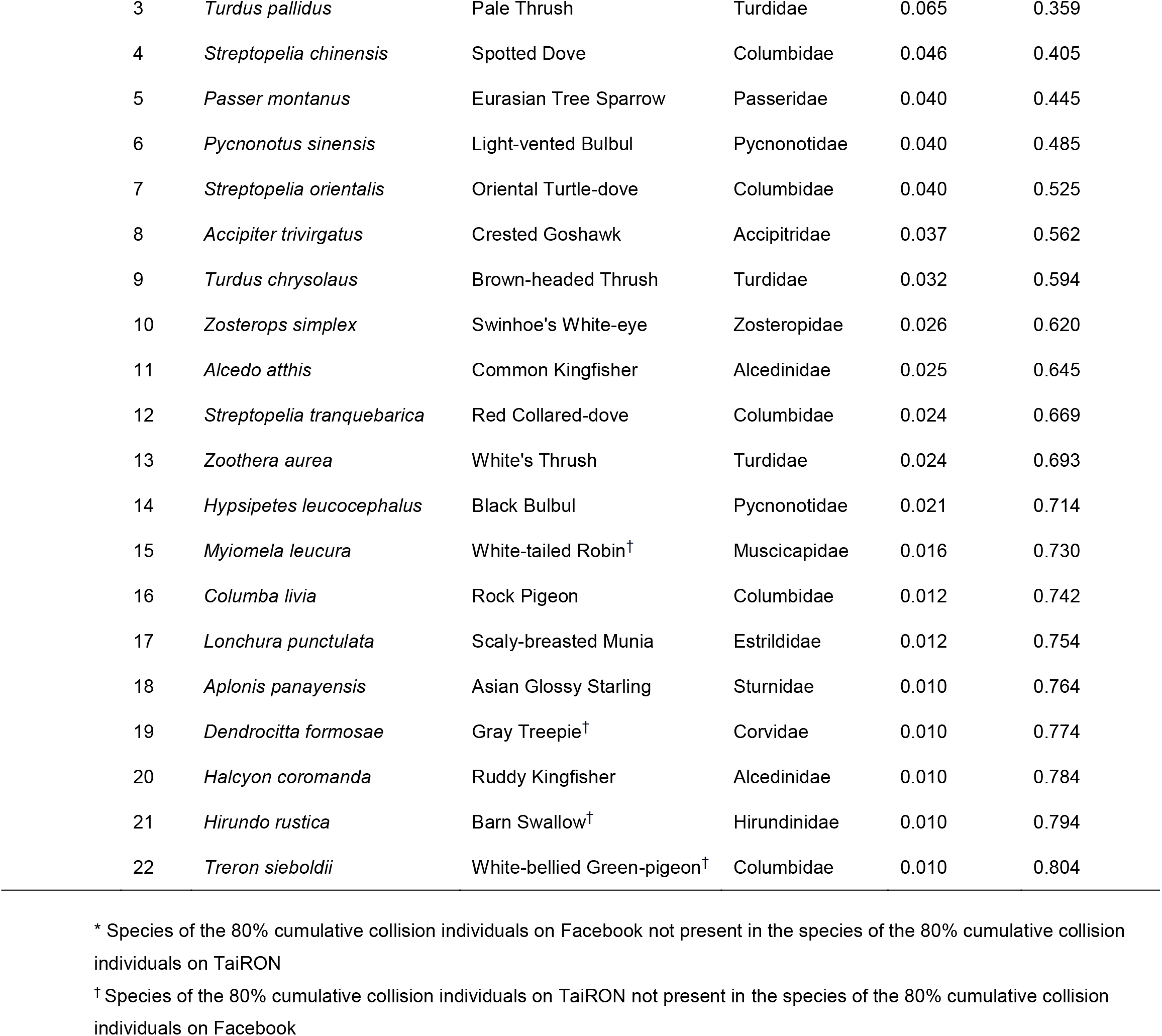
Species compositions of the 80% cumulative collision individuals in the data between 2012 and 2022 collected from Facebook and TaiRON. Note that individuals not identified to species were excluded from the calculation.

| Platform | Rank | Species | Common name | Family | Proportion | Cumulative proportion |
| --- | --- | --- | --- | --- | --- | --- |
| Facebook | 1 | *Psilopogon nuchalis* | Taiwan Barbet | Megalaimidae | 0.192 | 0.192 |
|  | 2 | *Chalcophaps indica* | Asian Emerald Dove | Columbidae | 0.092 | 0.284 |
|  | 3 | *Passer montanus* | Eurasian Tree Sparrow | Passeridae | 0.061 | 0.345 |
|  | 4 | *Accipiter trivirgatus* | Crested Goshawk | Accipitridae | 0.059 | 0.404 |
|  | 5 | *Alcedo atthis* | Common Kingfisher | Alcedinidae | 0.059 | 0.463 |
|  | 6 | *Streptopelia chinensis* | Spotted Dove | Columbidae | 0.058 | 0.521 |
|  | 7 | *Pycnonotus sinensis* | Light-vented Bulbul | Pycnonotidae | 0.051 | 0.572 |
|  | 8 | *Turdus pallidus* | Pale Thrush | Turdidae | 0.031 | 0.603 |
|  | 9 | *Zosterops simplex* | Swinhoe's White-eye | Zosteropidae | 0.029 | 0.632 |
|  | 10 | *Streptopelia orientalis* | Oriental Turtle-dove | Columbidae | 0.020 | 0.652 |
|  | 11 | *Streptopelia tranquebarica* | Red Collared-dove | Columbidae | 0.019 | 0.671 |
|  | 12 | *Zoothera aurea* | White's Thrush | Turdidae | 0.019 | 0.690 |
|  | 13 | *Halcyon coromanda* | Ruddy Kingfisher | Alcedinidae | 0.017 | 0.707 |
|  | 14 | *Hypsipetes leucocephalus* | Black Bulbul | Pycnonotidae | 0.016 | 0.723 |
|  | 15 | *Accipiter virgatus* | Besra* | Accipitridae | 0.015 | 0.738 |
|  | 16 | *Aplonis panayensis* | Asian Glossy Starling | Sturnidae | 0.014 | 0.752 |
|  | 17 | *Columba livia* | Rock Pigeon | Columbidae | 0.009 | 0.761 |
|  | 18 | *Lonchura punctulata* | Scaly-breasted Munia | Estrildidae | 0.009 | 0.770 |
|  | 19 | *Lanius cristatus* | Brown Shrike* | Laniidae | 0.008 | 0.778 |
|  | 20 | *Ninox japonica* | Northern Boobook* | Strigidae | 0.008 | 0.786 |
|  | 21 | *Turdus chrysolaus* | Brown-headed Thrush | Turdidae | 0.008 | 0.794 |
|  | 22 | *Cuculus optatus* | Oriental Cuckoo* | Cuculidae | 0.007 | 0.801 |
| TaiRON | 1 | *Psilopogon nuchalis* | Taiwan Barbet | Megalaimidae | 0.175 | 0.175 |
|  | 2 | *Chalcophaps indica* | Asian Emerald Dove | Columbidae | 0.119 | 0.294 |

|  |  |  |  |  |  |
| --- | --- | --- | --- | --- | --- |
| 3 | *Turdus pallidus* | Pale Thrush | Turdidae | 0.065 | 0.359 |
| 4 | *Streptopelia chinensis* | Spotted Dove | Columbidae | 0.046 | 0.405 |
| 5 | *Passer montanus* | Eurasian Tree Sparrow | Passeridae | 0.040 | 0.445 |
| 6 | *Pycnonotus sinensis* | Light-vented Bulbul | Pycnonotidae | 0.040 | 0.485 |
| 7 | *Streptopelia orientalis* | Oriental Turtle-dove | Columbidae | 0.040 | 0.525 |
| 8 | *Accipiter trivirgatus* | Crested Goshawk | Accipitridae | 0.037 | 0.562 |
| 9 | *Turdus chrysolaus* | Brown-headed Thrush | Turdidae | 0.032 | 0.594 |
| 10 | *Zosterops simplex* | Swinhoe's White-eye | Zosteropidae | 0.026 | 0.620 |
| 11 | *Alcedo atthis* | Common Kingfisher | Alcedinidae | 0.025 | 0.645 |
| 12 | *Streptopelia tranquebarica* | Red Collared-dove | Columbidae | 0.024 | 0.669 |
| 13 | *Zoothera aurea* | White's Thrush | Turdidae | 0.024 | 0.693 |
| 14 | *Hypsipetes leucocephalus* | Black Bulbul | Pycnonotidae | 0.021 | 0.714 |
| 15 | *Myiomela leucura* | White-tailed Robin<sup>†</sup> | Muscicapidae | 0.016 | 0.730 |
| 16 | *Columba livia* | Rock Pigeon | Columbidae | 0.012 | 0.742 |
| 17 | *Lonchura punctulata* | Scaly-breasted Munia | Estrildidae | 0.012 | 0.754 |
| 18 | *Aplonis panayensis* | Asian Glossy Starling | Sturnidae | 0.010 | 0.764 |
| 19 | *Dendrocitta formosae* | Gray Treepie<sup>†</sup> | Corvidae | 0.010 | 0.774 |
| 20 | *Halcyon coromanda* | Ruddy Kingfisher | Alcedinidae | 0.010 | 0.784 |
| 21 | *Hirundo rustica* | Barn Swallow<sup>†</sup> | Hirundinidae | 0.010 | 0.794 |
| 22 | *Treron sieboldii* | White-bellied Green-pigeon<sup>†</sup> | Columbidae | 0.010 | 0.804 |

\* Species of the 80% cumulative collision individuals on Facebook not present in the species of the 80% cumulative collision individuals on TaiRON † Species of the 80% cumulative collision individuals on TaiRON not present in the species of the 80% cumulative collision individuals on Facebook

Regarding the top BWC species that contributed to the 50% cumulative BWC individuals on the two platforms, the first two species were the same, being Taiwan Barbet (*Psilopogon nuchalis*) and Asian Emerald Dove (*Chalcophaps indica*). The remaining species were Eurasian Tree Sparrow (*Passer montanus*), Crested Goshawk (*Accipiter trivirgatus*), and Common Kingfisher (*Alcedo atthis*) on Facebook, and Pale Thrush (*Turdus pallidus*), Spotted Dove (*Streptopelia chinensis*), Eurasian Tree Sparrow, Light-vented Bulbul (*Pycnonotus sinensis*), and Oriental Turtle-dove (*Streptopelia orientalis*) on TaiRON (species were ordered by their rankings) (Table 1).

### 3.3 Sampling completeness of the two platforms

The observed sampling completeness of the BWC data from Facebook and TaiRON were 0.980 (95% CI: 0.975–0.984) and 0.960 (95% CI: 0.951–0.969), respectively (Fig. 7), confirming the adequacy of the data collection efforts. The number of BWC species and sampling completeness did not differ between Facebook and TaiRON at the rarefied reference point (number of BWC species: Facebook vs. TaiRON [95% CI] = 106 [100–112] vs. 104 [95–113]; sampling completeness: Facebook vs. TaiRON [95% CI] = 0.958 [0.953–0.963] vs. 0.960 [0.951–0.969]) (Fig. 7), suggesting that the data quality of the two platforms were comparable.

**Figure 7.**
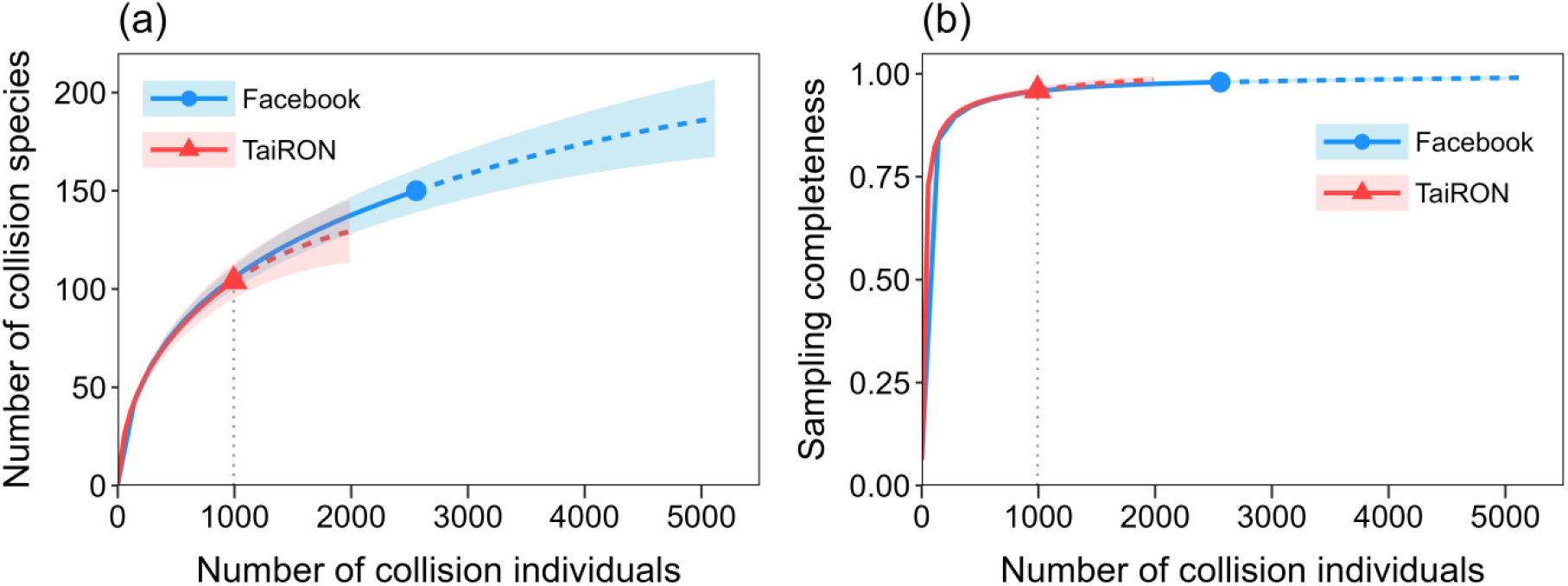
(a) Sample-size-based interpolation (solid lines) and extrapolation (dashed lines) sampling curves and (b) sampling completeness curves for the collision data between 2012 and 2022 collected from Facebook and TaiRON. Points indicate the observed values; shaded areas represent 95% confidence intervals of the estimates; gray dotted lines denote the rarefied reference points. Note that collision individuals not identified to the species level were excluded from the analysis.

## 4. Discussion

To our knowledge, this study is among the first to collect BWC data through social media posts. Compared to the number of BWC cases from the dedicated citizen science platform TaiRON, there were more than twice as many cases from Facebook, along with more information regarding the states of the BWC individuals. Moreover, the nationwide geographical coverage, the species compositions, and the sampling completeness of the BWC data were similar between the two platforms, suggesting that their data quality is comparable. BWC is an overlooked threat to birdlife in Asia, and with the assistance of Facebook citizen science efforts, we can obtain decent BWC records in terms of both quantity and quality.

The levels of public participation and engagement for citizen science projects on social media platforms are generally lower than those on dedicated platforms. According to the definitions in Haklay (2013), the Facebook group “Report on Bird-Glass Collisions” is classified as a citizen science project located between level one “Crowdsourcing” (citizens as sensors) and level two “Distributed Intelligence” (citizens as basic interpreters), whereas TaiRON is classified as a project between level two and level three “Participatory Science” (citizens as both issue proposers and data collectors). Despite such differences, our results show that the nationwide geographical coverage, the species compositions, and the sampling completeness of the BWC data from these two platforms were indeed comparable. This suggests that the roles of participants in citizen science projects may not necessarily affect the quality of the BWC data, and social media posts can provide a great source of citizen science observations.

The two platforms exhibited similar species rank-abundance distributions and species compositions of the top 80% cumulative BWC individuals. When both data sources were combined, the number of BWC species increased from 153 and 104 for Facebook and TaiRON respectively to 172 species (Fig. 5), demonstrating that integrating different citizen science data sources can help reveal a broader pool of BWC species, especially those low in BWC numbers yet globally threatened (e.g., Fairy Pitta *Pitta nympha*) or endemic with a small population (e.g., Swinhoe’s Pheasant *Lophura swinhoii)*. Furthermore, our data revealed distinct families of BWC species from those reported in North America, South America, and Europe (Klem Jr 1989, Loss et al. 2014, Basilio et al. 2020), with higher BWC numbers of Megalaimidae, Columbidae, Passeridae, Accipitridae, Alcedinidae, and Pycnonotidae. In fact, this is the first study to report a species in the Megalaimidae family, Taiwan Barbet, as the top super-collider, as well as a raptor species in the Accipitridae family, Crested Goshawk, as a super-collider. These findings highlight the need for more BWC studies in Asia to develop appropriate conservation strategies.

There are several potential advantages of collecting BWC data on the social media Facebook compared to the dedicated platform TaiRON. First, Facebook is more accessible to the public. TaiRON users are trained to report their observations in standardized entry formats, whereas Facebook users can post any contents for their findings in any formats, facilitating reporting observations by those unfamiliar with the reporting procedures on TaiRON. Second, there are more users on Facebook than on TaiRON, allowing for a wider search for BWC birds (e.g., more than twice cases from Facebook, with more than three times species present only on Facebook than only on TaiRON [Fig. 5]). A combination of active posting and passive crowdsourcing on social media (Ghermandi and Sinclair 2019) can help gather more BWC data than do the traditional citizen science platforms that rely mainly on active reporting. Third, BWC observations can be traced back to early years before the advent of dedicated citizen science platforms. For instance, by searching through Facebook posts, we were able to record a BWC case occurring more than three decades ago—a Tawny Fish-owl (*Ketupa flavipes*) that collided with glass panes at Hsinchu Train Station in 1992.

Immediate posting by Facebook users can provide important information regarding the states of the BWC individuals. In our Facebook data, over half of the BWC individuals were non-dead when observed, which would not have otherwise been reported to TaiRON (Fig. 4). These “hidden BWCs” may explain why there were more than twice as many BWC cases on Facebook as those on TaiRON. Moreover, the project manager can educate and guide the observers to handle the BWC birds, for example, sending the injured individuals to rescue centers in an appropriate manner and the dead individuals to museums for long-term collections and further research. Thus, Facebook may serve as a bilateral platform where contributors can report BWC cases and meanwhile the project manager can communicate with the contributors. This helps promote public awareness and participation of young developing topics.

Despite the advantages of Facebook as a citizen science data source, several limitations exist. Some information such as the GPS coordinates of the BWC cases may be incomplete as the contributors might not provide the exact locations of their observations. Moreover, because most Facebook contributors may not have enough background knowledge in bird species identification, their reports need to be double-checked to ensure the accuracy. Finally, since it is not possible to automatically retrieve BWC data from Facebook because of the variable contents and formats, searching for relevant posts, requesting detailed information, and recording the data can only be done manually. Nonetheless, even though the data collection process is labor-demanding and time-consuming, the quantity and quality of the resulting BWC data may still outweigh the costs.

It is generally believed that citizen science data from social media may be of lower quality and thus are not as valuable as those collected from dedicated platforms. However, our study shows that the quantity and quality of BWC data collected from Facebook can indeed be comparable to those from the well-developed citizen science platform TaiRON. Furthermore, Facebook provides greater opportunities to disseminate knowledge and educational contents about BWCs. Taken together, Facebook has a great potential not only as a tool for citizen science projects, but also as an education platform for the public. Finally, integrating data from different citizen science sources can help paint a more complete picture of BWC patterns that benefits conservation management.

## Acknowledgments

We thank Te-En Lin for providing BWC data reported to TaiRON, Chia-Yun Kan for assisting with collecting and arranging BWC cases on Facebook, and Jian-Long Wu, Cheng-Te Yao, and Yen-Hsing Lin for assisting with identifying BWC species. We also thank all the citizens who have provided their BWC observations on the Facebook group “Reports on Bird-Glass Collisions” and TaiRON.

## Author contributions

C.-H. Hsieh and L.-M. Wang conceived the ideas; C.-H. Hsieh and L.-M. Wang collected the data; C.-H. Hsieh and G.-C. Hsu analyzed the data; C.-H. Hsieh and G.-C. Hsu wrote the first draft of the manuscript. All authors revised the manuscript and approved the final version for publication.

## Data availability statement

Data and code used in this manuscript will be publicly available on Zenodo if the manuscript is accecpted for publication.

