## Appendix Methods for "Can social media serve as a potential citizen science source for bird-window collision (BWC) data? A study using a decadal data set in Taiwan"

Appendix Methods: The search queries for collecting bird-window collision data from public Facebook posts.

Between February 2020 and December 2022, we carried out systematic searches through public Facebook posts on a weekly basis using a series of designed search terms as listed in the table below. The search was conducted in Taiwanese Mandarin, and the English translation for the search terms were provided.

| Search term (Taiwanese Mandarin) | Search term (English translation) |
|---|---|
| Not able to provide | bird & collide |
| Not able to provide | bird & collide with glass |
| Not able to provide | bird & collide with window |
| Not able to provide | bird & collide with door |
| Not able to provide | bird & collide with wall |

People tend to describe certain families of birds by their common names if they were able to identify the collision individuals. In case the users used the common names instead of the word "bird" *per se* in their posts, we conducted additional search each month by replacing the keyword "bird" with the common names listed in the table below.

| Common name (Taiwanese Mandarin) | Corresponding bird family (English translation) |
|---|---|
| Not able to provide | Accipitriformes |
| Not able to provide | Anseriformes |
| Not able to provide | Galliformes |
| Not able to provide | Passeriformes |
| Not able to provide | Strigiformes |
